# A Pseudo-Longitudinal Methylome Projection Framework Defines a Buccal PACE-like Aging-Rate Score from Cross-Sectional DNA Methylation Data

**DOI:** 10.64898/2026.08.03.742627

**Authors:** Tatsuma Shoji, Ryo Nakaki

**Author notes:** **Correspondence:** Tatsuma Shoji.

## Abstract

**Background:** DNA methylation-based biomarkers have enabled robust estimation of biological age across tissues, and longitudinally trained measures such as DunedinPACE provide estimates of the pace of aging from blood methylomes. However, longitudinal methylation data are often unavailable, particularly for minimally invasive tissues such as buccal mucosa. Here, we developed a pseudo-longitudinal framework to estimate a buccal mucosa-derived PACE-like aging-rate score from cross-sectional methylome data.

**Methods:** We used a buccal biological age estimator as an internal pseudo-time axis. Methylation beta-values were transformed to M-values, and CpG-specific smooth functions of biological age were fitted in cross-validation. Local derivatives of these functions were used to project each individual’s buccal methylome forward by a small time step. The projected methylome was converted back to beta-values, biological age was recalculated, and the change in biological age per unit time was defined as a pseudo-aging velocity. This raw velocity was transformed to a non-negative PACE-like score centered at 1.0. We then trained cross-fitted models to predict the derived score from buccal CpG methylation profiles.

**Results:** In 151 individuals, the proposed score was reproducibly predicted from buccal methylomes in out-of-fold analysis, with a Pearson correlation of 0.706 and Spearman correlation of 0.710 between observed and predicted PACE-like scores. Sensitivity analyses across CpG selection size and regression models showed broadly consistent performance. In contrast, the proposed buccal PACE-like score showed only modest association with measured DunedinPACE, and alternative attempts to reconstruct DunedinPACE from buccal methylomes, including supervised proxy modeling and buccal-to-blood CpG imputation, showed limited sample-level performance.

**Conclusions:** These results support the feasibility of deriving a tissue-specific PACE-like aging-rate score from cross-sectional buccal methylome data by treating biological age as a pseudo-time axis. The proposed score should not be interpreted as a replacement for blood-derived DunedinPACE, but rather as an exploratory buccal methylome dynamics index that may capture tissue-specific aging-related variation.

## 1. Introduction

DNA methylation has emerged as one of the most reproducible molecular readouts of human aging. Early DNA methylation clocks demonstrated that chronological age can be accurately estimated from methylation levels at selected CpG sites, both across multiple tissues and within blood-derived datasets [1,2]. Subsequent biomarkers, including DNAm PhenoAge and DNAm GrimAge, extended this concept by training methylation predictors on phenotypic age, mortality-related proteins, smoking surrogates, and healthspan-related outcomes [3,4]. These developments have shifted the field from estimating chronological age toward quantifying biological aging and disease-relevant aging heterogeneity [5,6].

Among current methylation-based aging biomarkers, DunedinPACE is particularly relevant because it was developed to approximate the longitudinal pace of aging. It was trained using blood DNA methylation to predict a longitudinal Pace of Aging phenotype derived from within-person changes in multiple organ-system biomarkers in the Dunedin Study. DunedinPACE has been reported to show test-retest reliability and associations with morbidity, disability, mortality, and early-life adversity [7]. However, DunedinPACE is fundamentally a blood-derived biomarker trained against longitudinal phenotypic change, and its direct application to other tissues, such as buccal mucosa, may not capture identical biological information.

Buccal mucosa is attractive for aging biomarker development because it can be collected non-invasively and repeatedly. However, most buccal methylome datasets are cross-sectional, limiting the direct estimation of individual aging rates. This creates a methodological gap: although longitudinal data are ideal for estimating rates of biological change, many practical cohorts contain only one methylation profile per individual. A method that extracts pseudo-longitudinal information from cross-sectional buccal methylomes would therefore be useful, particularly as a tissue-specific exploratory approach rather than as a direct substitute for blood-trained pace-of-aging biomarkers.

The idea of ordering high-dimensional molecular profiles along an inferred progression axis is well established in single-cell transcriptomics. Pseudotime algorithms, such as Monocle, infer cellular trajectories from cross-sectional single-cell data and have been used to reconstruct dynamic biological processes from static molecular snapshots [8]. Although individuals in a methylome cohort are not cells within a differentiation process, a related concept can be adapted: if a biologically meaningful ordering axis is available, local methylation changes along that axis may approximate pseudo-longitudinal molecular dynamics.

Here, we propose a pseudo-longitudinal methylome projection framework for buccal DNA methylation data. Instead of attempting to infer chronological time directly, we use a buccal biological age estimator as an internal pseudo-time axis. For each CpG, methylation M-values are modeled as smooth functions of biological age, local derivatives are computed, and each individual’s methylome is projected forward by a small time step. Biological age is then recalculated from the projected methylome, and the resulting change is converted into a non-negative PACE-like score centered at 1.0. We further evaluate whether this score can be predicted from buccal methylomes in out-of-fold analysis and whether it overlaps with blood-derived DunedinPACE or with direct buccal-to-blood DunedinPACE reconstruction strategies.

## 2. Results

### 2.1. Direct supervised prediction of DunedinPACE from buccal methylomes showed limited performance

Before applying the proposed pseudo-longitudinal framework, we first asked whether blood-derived DunedinPACE could be reconstructed directly from buccal methylomes. Measured DunedinPACE was used as the target variable, and buccal DNA methylation profiles were used as predictors in an out-of-fold ElasticNet model. As shown in Figure 1, the ElasticNet DunedinPACE proxy achieved only weak out-of-fold performance, with Pearson r = 0.216 and Spearman ρ = 0.201. The predicted values were substantially compressed toward the mean, indicating limited recoverable DunedinPACE signal from buccal methylation profiles in this cohort. These results argue against the idea that a simple supervised model can robustly reconstruct measured DunedinPACE from buccal methylomes and motivated the development of a tissue-specific buccal PACE-like score rather than a buccal implementation of DunedinPACE.

**Figure 1.**
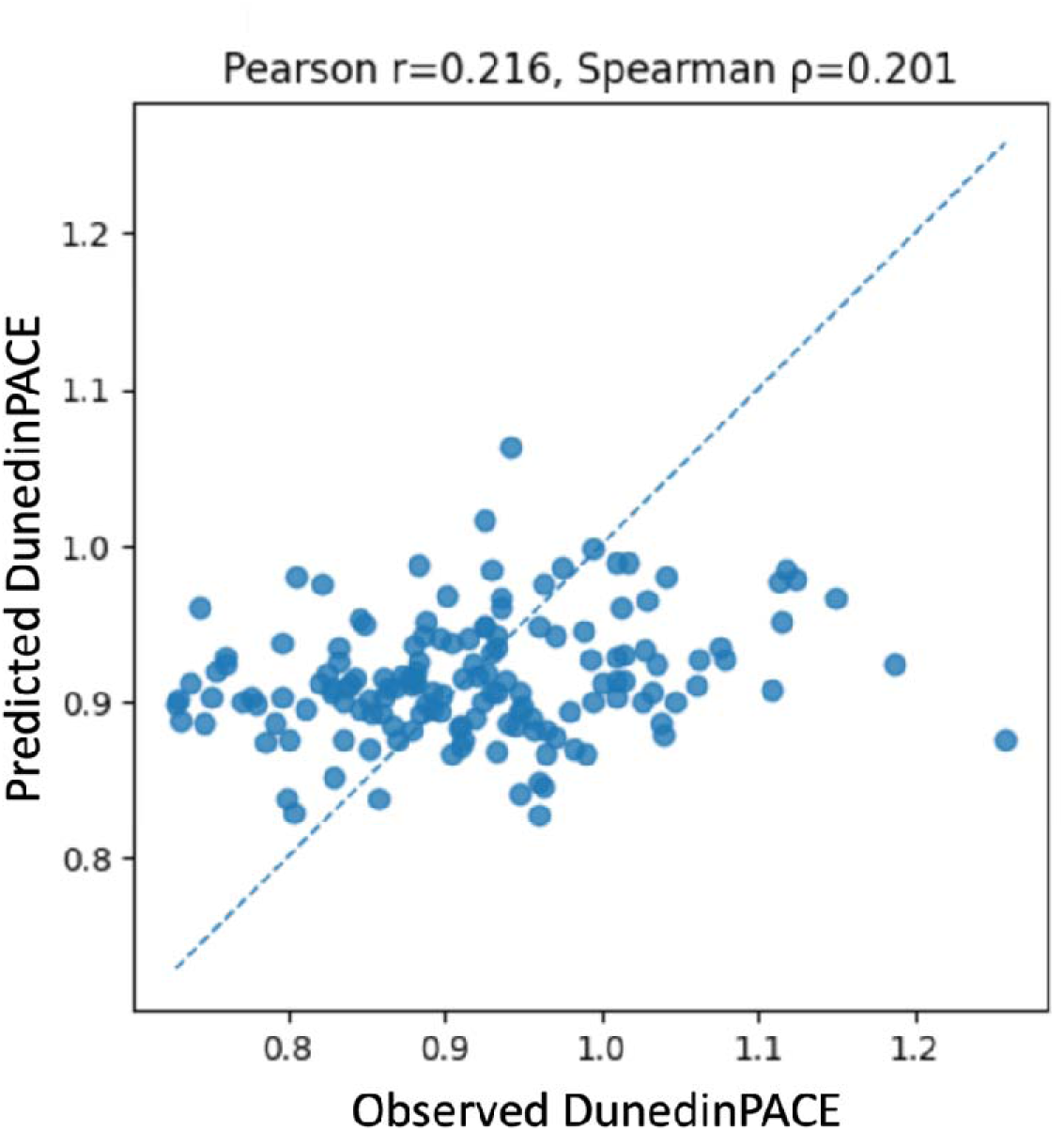
Direct ElasticNet prediction of DunedinPACE from buccal methylomes. Measured DunedinPACE was used as the target variable, and buccal DNA methylation profiles were used as predictors in an out-of-fold ElasticNet model. Each point represents one participant. The dashed line indicates the identity line. The predicted values were compressed toward the mean, indicating limited recoverable DunedinPACE signal from buccal methylomes in this cohort. Pearson r = 0.216 and Spearman ρ = 0.201.

### 2.2. Buccal-to-blood CpG imputation captured global CpG-level methylation structure but did not adequately reconstruct sample-level DunedinPACE

As a second preliminary strategy, we further explored whether reconstructing blood methylation values at DunedinPACE-related CpGs from buccal methylation could improve DunedinPACE prediction. At the pooled CpG-pair level, imputed blood beta-values showed a high overall correlation with observed blood beta-values across all subjects and target CpGs, with Pearson r = 0.984 and Spearman ρ = 0.983 across 25,670 observed-predicted CpG pairs (Figure 2). However, this high pooled correlation primarily reflects recovery of broad CpG-level methylation distributions and average methylation levels across CpGs. It did not translate into strong sample-level DunedinPACE reconstruction: DunedinPACE calculated from imputed blood CpGs showed only modest correlation with observed DunedinPACE, with Pearson r = 0.211 and Spearman ρ = 0.247 (Figure 3). This distinction is important. A model may accurately reproduce the global distribution of methylation values across CpGs while still failing to reconstruct the weighted within-sample pattern required for a composite biomarker such as DunedinPACE. Thus, buccal-to-blood CpG imputation did not provide a superior route for recovering blood-derived DunedinPACE in this dataset.

**Figure 2.**
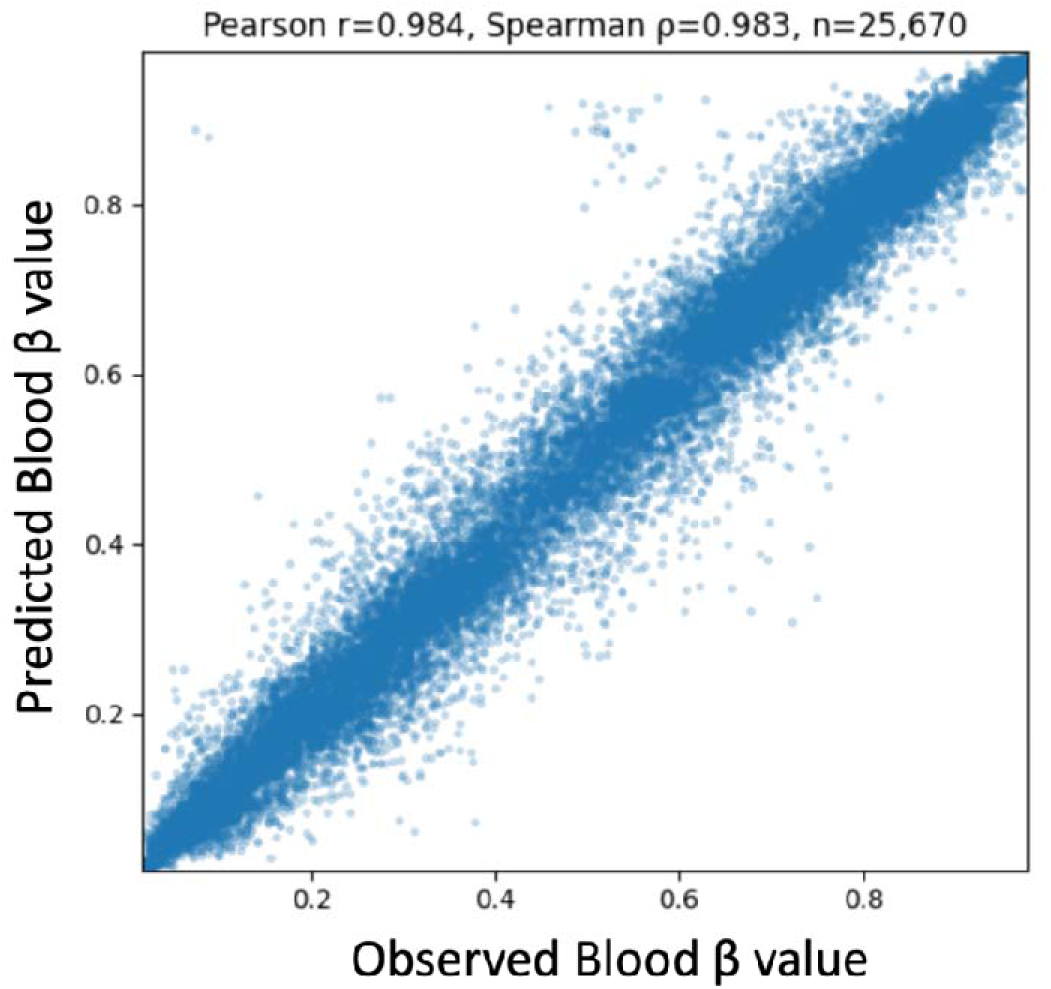
Pooled CpG-level association between observed and imputed blood beta-values. Blood beta-values at DunedinPACE-related CpG sites were predicted from buccal methylation profiles, and the observed and imputed blood beta-values were pooled across all participants and target CpGs. Each point represents one subject–CpG pair. The high pooled correlation indicates that the imputation model recovered broad CpG-level methylation distributions and average methylation patterns across CpGs. However, this pooled CpG-level agreement should not be interpreted as accurate reconstruction of sample-level DunedinPACE. Pearson r = 0.984, Spearman ρ = 0.983, n = 25,670 observed–predicted CpG pairs.

**Figure 3.**
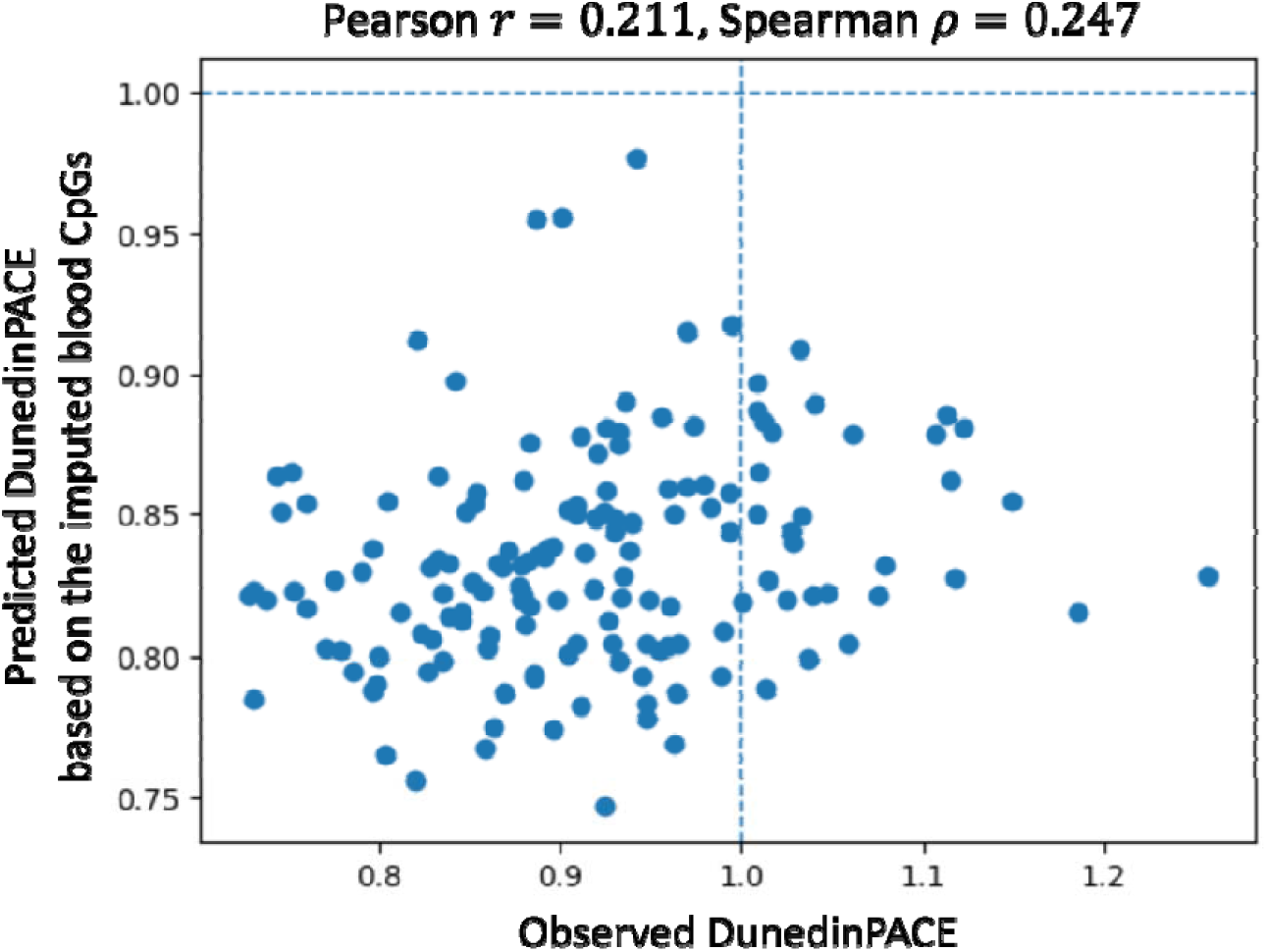
DunedinPACE calculated from imputed blood CpGs predicted from buccal methylomes. Blood methylation values at DunedinPACE-related CpGs were first imputed from buccal methylation profiles, and DunedinPACE was then recalculated using the imputed blood CpG matrix. Each point represents one participant. Dashed horizontal and vertical lines indicate a value of 1.0. Despite the high pooled CpG-level agreement shown in Figure 2, the imputed CpG matrix showed only modest performance for reconstructing individual-level DunedinPACE, suggesting that recovering global CpG methylation distributions is insufficient to reproduce the weighted within-sample methylation pattern required for DunedinPACE. Pearson r = 0.211 and Spearman ρ = 0.247.

### 2.3. Pseudo-longitudinal methylome projection framework and reproducible prediction of buccal PACE-like scores

Having established that direct supervised reconstruction of blood-derived DunedinPACE and buccal-to-blood CpG imputation were insufficient for robust sample-level DunedinPACE recovery, we next developed a tissue-specific buccal PACE-like score. The proposed framework is summarized in Figure 4. Starting from a cross-sectional buccal methylation beta-value matrix, beta-values were transformed to M-values to improve statistical modeling properties [9]. A pre-specified buccal biological age estimator was then used to assign each participant a biological age value, which was treated as an internal pseudo-time coordinate. For each selected CpG, M-values were modeled as smooth functions of biological age using spline-based regression. The local derivative of each CpG-specific function was evaluated at each individual’s biological age, providing an estimated methylation-change vector in high-dimensional methylome space. This derivative vector was used to project the individual’s current methylome forward by a small time step. The projected M-values were converted back to beta-values, biological age was recalculated, and the change in biological age per unit time was defined as the raw pseudo-aging velocity. Finally, this raw velocity was transformed to a non-negative PACE-like score with mean approximately 1.0. This workflow was designed to emulate, in a pseudo-longitudinal manner, the conceptual structure of an aging-rate measure: individuals with values above 1.0 are interpreted as showing faster projected biological-age advancement along the inferred buccal methylome aging axis, whereas individuals below 1.0 are interpreted as showing slower projected advancement. Importantly, the score is not a direct longitudinal aging rate, but a cross-sectional surrogate derived from local methylome dynamics along a biological-age axis.

**Figure 4.**
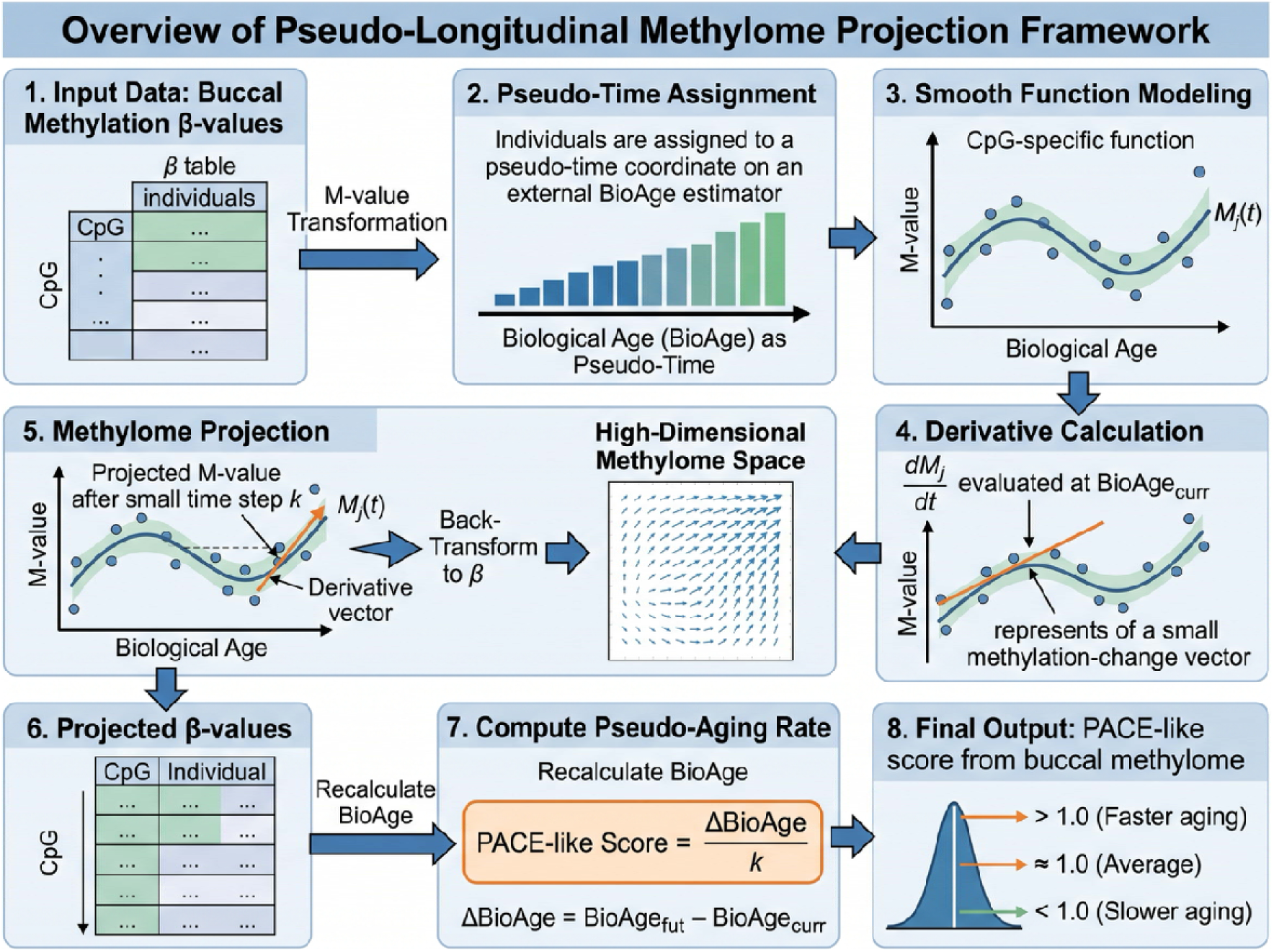
Conceptual overview of the pseudo-longitudinal buccal methylome projection framework. Cross-sectional buccal methylation beta-values are transformed to M-values. A buccal biological age model is used to assign each participant a pseudo-time coordinate. For selected CpGs, M-values are modeled as smooth functions of biological age, and local derivatives are estimated at each participant’s biological age. Each methylome is projected forward by a small time step along the inferred derivative field. Projected M-values are converted back to beta-values, biological age is recalculated, and the biological age change per unit time is defined as a raw pseudo-aging velocity. The raw velocity is transformed to a non-negative PACE-like score centered at 1.0.

Using this framework, the primary cross-fitted analysis demonstrated that the proposed buccal PACE-like score could be predicted from buccal methylome data with substantial out-of-fold accuracy. As shown in Figure 5, the correlation between observed and predicted PACE-like scores was Pearson r = 0.706 and Spearman ρ = 0.710 with k = 1 and 500 CpGs used for methylome dynamics estimation. The predicted values tracked the observed score across the full score range, although moderate shrinkage toward the mean was observed, as expected for regularized high-dimensional prediction. This result supports the internal reproducibility of the derived pseudo-aging velocity. In other words, although the score itself is generated from a pseudo-longitudinal projection procedure, the resulting score is not merely numerical noise: it can be recovered from independent held-out methylation profiles in cross-validation.

**Figure 5.**
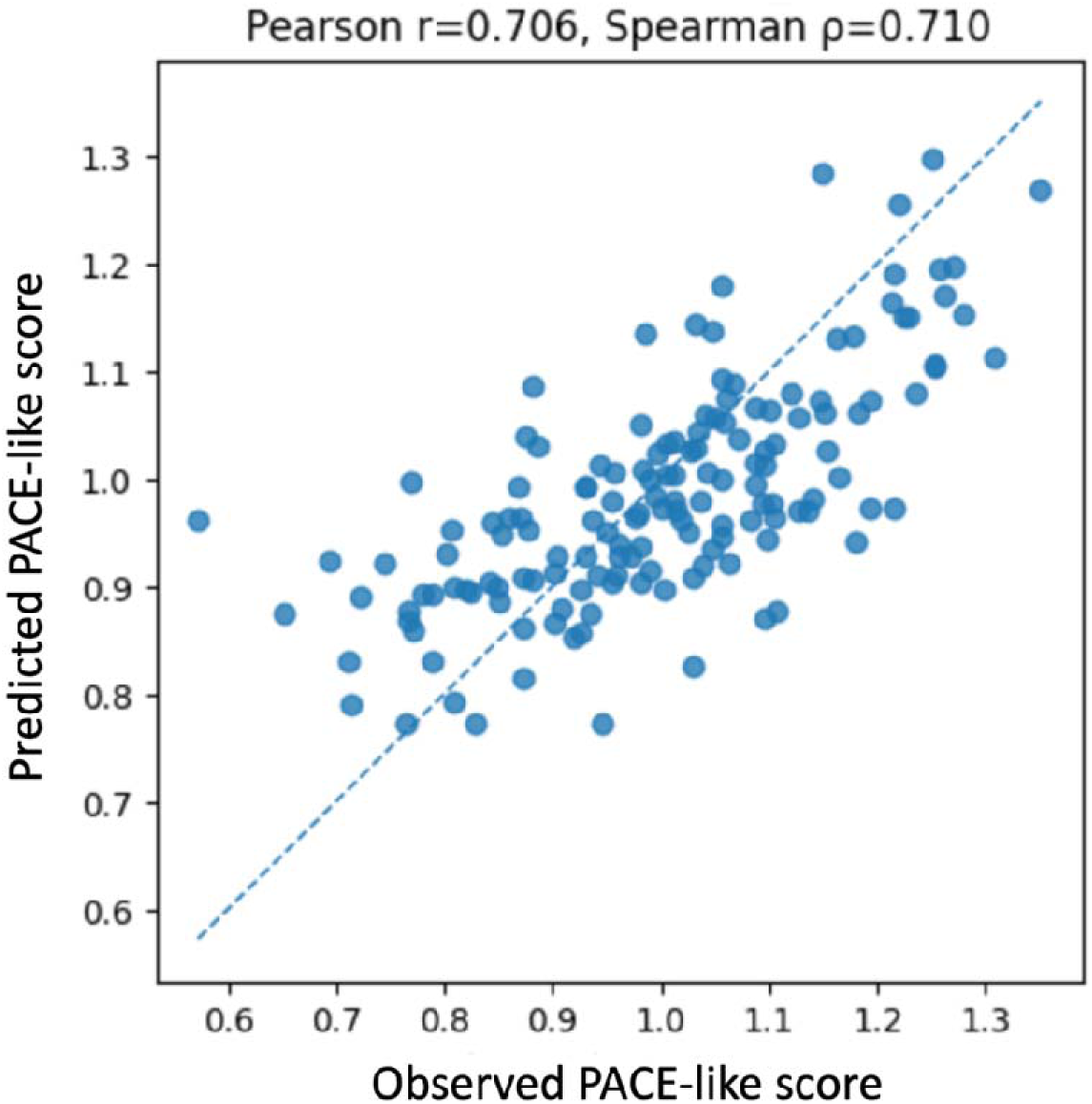
Out-of-fold prediction of the buccal PACE-like score. Scatter plot of observed versus predicted PACE-like scores in out-of-fold cross-validation. The predicted score was obtained for each participant using models trained without that participant. The dashed line indicates the identity line. Pearson r = 0.706 and Spearman ρ = 0.710 correlations are shown in the plot.

### 2.4. Sensitivity analysis across projection and prediction parameters

We then examined whether the observed performance was robust to key modeling choices, including the number of CpGs used for methylome dynamics estimation and the regression model used for predicting the final PACE-like score from buccal CpGs. The out-of-fold Pearson correlations between observed and predicted PACE-like scores ranged from 0.509 to 0.706 across the evaluated configurations (Table 1). The primary configuration, using k = 1, 500 dynamic CpGs, and Ridge regression, achieved the highest Pearson correlation of 0.706 and a Spearman correlation of 0.710. Ridge regression generally provided stable performance across CpG selection sizes, whereas ElasticNet performance was more variable, particularly when larger numbers of dynamic CpGs were used. These sensitivity analyses indicate that the method is not dependent on a single fragile hyperparameter setting. However, performance declined when very large numbers of dynamic CpGs were used with ElasticNet, consistent with the expectation that sparse feature selection may become unstable when the number of predictors greatly exceeds the number of participants.

**Table 1.**
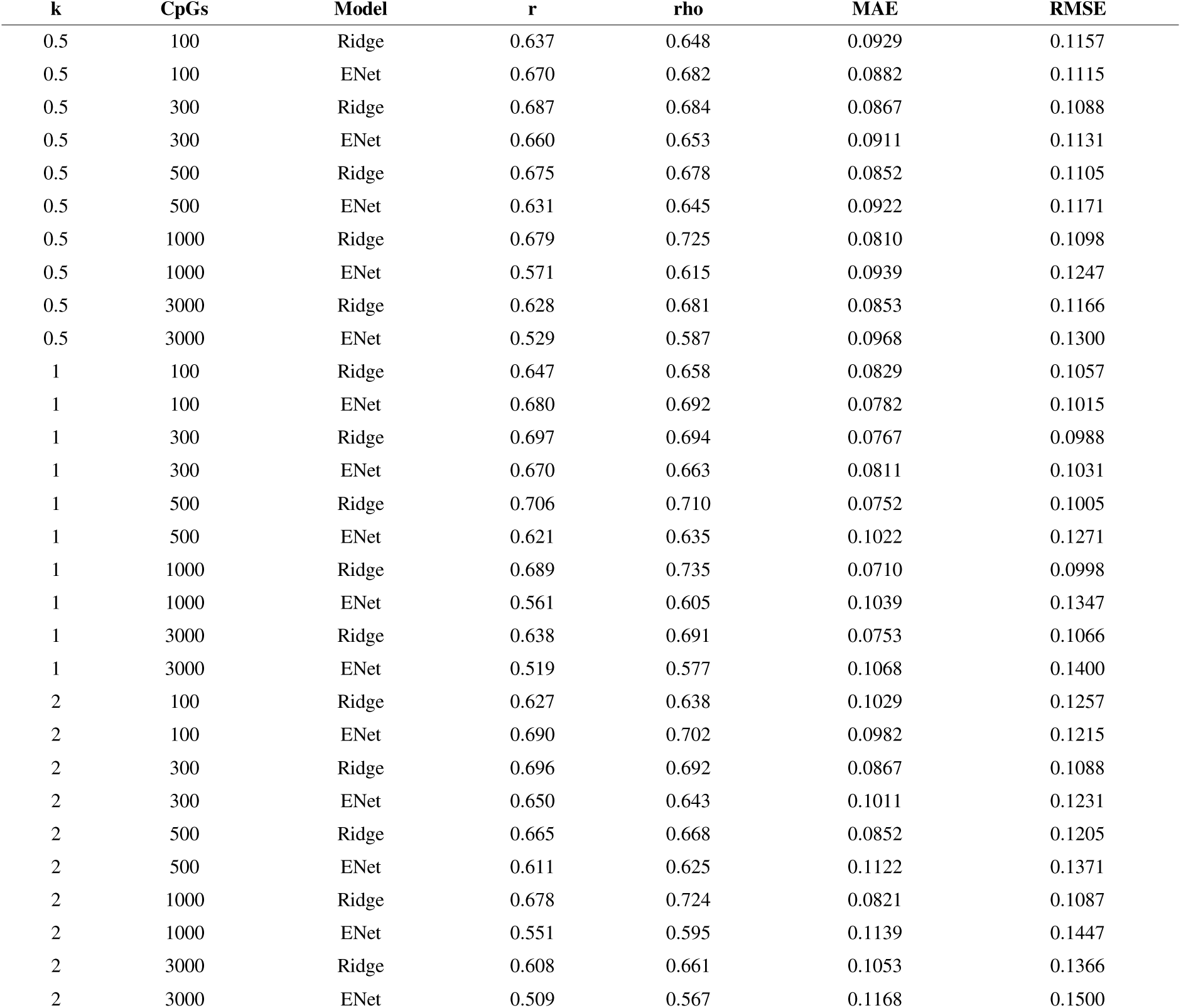
Sensitivity of observed-predicted PACE-like.

### 2.5. The buccal PACE-like score showed only modest association with measured DunedinPACE

Finally, we asked whether the proposed buccal PACE-like score recapitulates measured DunedinPACE. This comparison is important because DunedinPACE is an established pace-of-aging biomarker, but it is blood-derived and was trained on longitudinal physiological decline rather than on buccal methylome dynamics. In Figure 6, the observed buccal PACE-like score showed a modest correlation with DunedinPACE, with Pearson r = 0.281 and Spearman ρ = 0.301. This result suggests that the proposed score and DunedinPACE share limited common information, but are not interchangeable. The modest correlation is consistent with the interpretation that the proposed score captures a buccal methylome-specific pseudo-aging velocity rather than directly reconstructing the blood-derived DunedinPACE biomarker.

**Figure 6.**
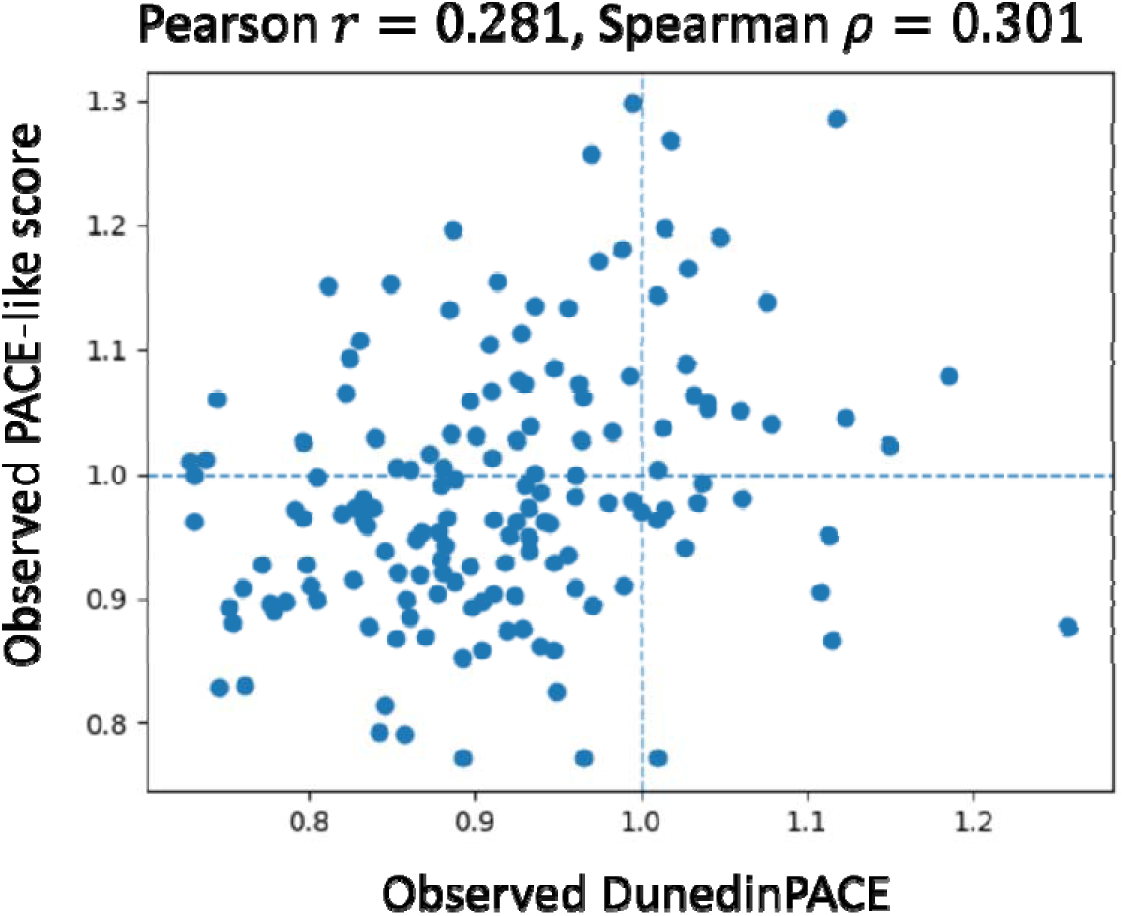
Association between measured DunedinPACE and the observed buccal PACE-like score. Each point represents one participant. The x-axis shows measured DunedinPACE, and the y-axis shows the observed buccal PACE-like score generated by the pseudo-longitudinal methylome projection framework. Dashed horizontal and vertical lines indicate a value of 1.0, corresponding to the conventional reference value for a pace-of-aging score. The modest association between the two measures indicates that the proposed buccal PACE-like score shares limited information with DunedinPACE but does not directly reproduce the blood-derived DunedinPACE biomarker. Pearson r = 0.281 and Spearman ρ = 0.301.

## 3. Methods

### 3.1. Study participants and sample collection

Whole blood and cheek mucosa samples were collected from healthy adult volunteers under an approved observational study protocol. A total of 151 unique participants were enrolled, and paired cheek mucosa and whole blood samples were obtained at Y’s Science Clinic Hiroo (Minato-ku, Tokyo, Japan) as part of the project titled “Evaluation of Biological Age Based on DNA Methylation and Its Clinical Significance,” reviewed by the Institutional Review Board of Shiba Palace Clinic. Written informed consent was obtained from all participants before sample collection. Genomic DNA was extracted from whole blood using the Maxwell RSC Blood DNA Kit (Promega, Madison, WI, USA) following the manufacturer’s instructions. Genomic DNA from cheek mucosa samples was extracted using Maxwell RSV Stabilized Saliva DNA Kit (Promega, Madison, WI, USA) following the manufacturer’s instructions. All procedures were conducted in accordance with the Ethical Guidelines for Medical and Biological Research Involving Human Subjects issued by the Japanese government.

### 3.2. DNA Methylation Profiling and Preprocessing

Genome-wide DNA methylation profiling was performed in a manner similar to that described by Shoji et al. [10]. Genomic DNA derived from both cheek mucosa and whole blood samples was analyzed using the Illumina Infinium HumanMethylationEPIC v2 BeadChip (EPICv2). Rhelixa, Inc. (Chuo-ku, Tokyo, Japan) processed and scanned the array. Following bisulfite conversion according to the manufacturer’s recommended protocol, the converted DNA was amplified, fragmented, hybridized to the arrays, and scanned using an Illumina iScan System. Raw IDAT files were processed in R (version 4.4.2) using the SeSAMe pipeline (version 1.24.0) [11, 12], including background correction with the normal-exponential out-of-band (noob) method and probe-level quality filtering using the pOOBAH detection mask. DNA methylation beta values were calculated as the ratio of methylated signal intensity to the sum of methylated and unmethylated signal intensities [13]. Before downstream analyses, probes that failed detection thresholds, those overlapping with single-nucleotide polymorphisms likely to affect hybridization, those exhibiting cross-reactive mapping to multiple genomic loci, or those located on sex chromosomes were excluded. Raw IDAT files and processed beta value matrices are available upon reasonable request.

### 3.3. Beta-value to M-value transformation

Methylation beta-values were transformed to M-values before spline modeling of CpG dynamics:

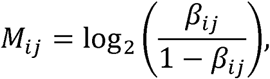

where *β_ij_* is the methylation beta-value of CpG*_j_* in participant *i*. Beta-values were clipped to a small interval away from 0 and 1 before transformation to avoid infinite values. The M-value scale was used for modeling because it is generally more statistically appropriate for differential or regression-based methylation analyses, whereas beta-values are more directly interpretable biologically [9].

### 3.4. Biological age estimation

Biological age was calculated using a pre-specified buccal methylation age model, referred to here as EpiclockAgeMB1 [14]. The model was applied as a linear predictor:

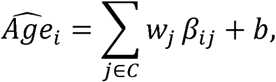

Where *C* is the set of CpGs included in the biological age model, *w_j_* is the corresponding coefficient, and *b* = 51.024577 is the intercept. This biological age estimate was used as the pseudo-time coordinate for the main Method 1 analysis. DunedinPACE estimation was performed according to the model introduced by Belsky et al. [7].

### 3.5. CpG selection for methylome dynamics estimation

To reduce computational burden and improve stability, CpGs used for dynamic modeling were selected within each training fold. Candidate CpGs were restricted to those present in both the methylation matrix and the biological age model feature set. CpGs with insufficient non-missingness or near-zero variance were excluded. Among remaining CpGs, the top CpGs most strongly associated with biological age were selected. The number of selected CpGs was varied in sensitivity analyses.

### 3.6. Spline-based modeling of CpG methylation as a function of biological age

For each selected CpG, M-values were modeled as a smooth function of biological age:

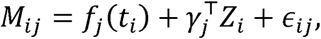

where *M_ij_* is the M-value for participant *i* and CpG *j*, *t_i_* is the estimated biological age, *Z_i_* represents optional covariates such as sex, and *f_j_* is a spline function. In implementation, multi-output ridge regression was used with a spline basis expansion of biological age. This enabled simultaneous fitting of CpG-specific smooth functions while controlling model complexity.

### 3.7. Projection of methylomes into a pseudo-future state

The local derivative of each fitted CpG function was computed with respect to biological age:

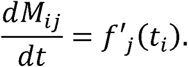

For a projection interval *k*, the pseudo-future M-value was defined as:

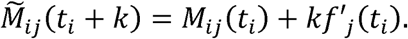

In the primary analysis, *k* = 1 was used to represent a one-year step along the biological-age axis. The projected M-values were transformed back to beta-values:

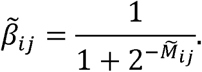

### 3.8. Definition of raw pseudo-aging velocity and PACE-like score

Biological age was recalculated from the projected beta-value matrix. The raw pseudo-aging velocity was defined as:

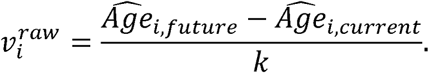

The raw velocity was then transformed into a non-negative PACE-like score centered at 1.0. Specifically, raw values were centered and exponentiated:

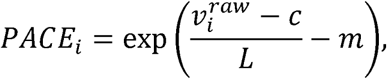

where *c* is the mean of the raw velocity in the training set, *L* is a scale parameter selected to target an approximate standard deviation of 0.15, and *m* is a normalization constant ensuring that the mean score in the training set is approximately 1.0. This transformation produces positive values with an intuitive interpretation: values above 1.0 represent faster projected aging and values below 1.0 represent slower projected aging.

### 3.9. Cross-fitted prediction of the PACE-like score from buccal methylomes

To evaluate whether the derived PACE-like score could be recovered from buccal methylomes, we trained prediction models using the methylation matrix as input and the derived raw velocity as the target. The procedure was performed in an outer cross-fitting framework. In each outer fold, CpG selection, methylome dynamics modeling, PACE-like target derivation, scaling, and predictor training were conducted using only the training samples. The held-out samples were then scored using models trained without those samples. Ridge regression and ElasticNet were compared as prediction models. Within each training fold, CpGs were prefiltered by association with the target velocity to reduce dimensionality. Model hyperparameters were selected by inner cross-validation. The final out-of-fold predicted score was obtained for all participants by aggregating predictions across outer folds.

### 3.10. Sensitivity analyses

Sensitivity analyses evaluated the robustness of observed-predicted PACE-like agreement across projection and modeling choices. The number of dynamic CpGs was varied, and Ridge regression was compared with ElasticNet. The primary performance metric was the out-of-fold Pearson correlation between observed and predicted PACE-like scores. Spearman correlation, root mean squared error, and mean absolute error were also recorded.

### 3.11. Comparison with DunedinPACE and alternative reconstruction strategies

Blood-derived measured DunedinPACE was used as an external comparator. Three comparisons were performed. First, observed and predicted buccal PACE-like scores were correlated with blood-derived measured DunedinPACE. Second, blood-derived DunedinPACE was directly predicted from buccal methylomes using supervised regularized regression. Third, blood methylation values at blood-derived DunedinPACE-related CpGs were predicted from buccal methylomes, and DunedinPACE was recalculated from the imputed blood CpG matrix. The latter approach was evaluated both at the pooled CpG-pair level and at the sample-level DunedinPACE score level.

## 4. Discussion

In this study, we developed a pseudo-longitudinal methylome projection framework to derive a buccal mucosa PACE-like aging-rate score from cross-sectional DNA methylation data. The central idea is to use biological age as an internal pseudo-time coordinate, estimate local CpG methylation derivatives along that coordinate, project each participant’s methylome into a pseudo-future state, and recalculate biological age from the projected methylome. The resulting pseudo-aging velocity was transformed into a positive PACE-like score centered at 1.0. The main finding was that this derived score could be robustly predicted from buccal methylomes in out-of-fold analysis, with Pearson r = 0.706 and Spearman ρ = 0.710 between observed and predicted PACE-like scores.

This result supports the feasibility of estimating an aging-rate-like quantity from buccal methylome data even in the absence of longitudinal sampling. The proposed framework does not claim to observe true within-person aging trajectories. Instead, it constructs a pseudo-longitudinal local derivative field using cross-sectional variation ordered by biological age. This distinction is important. Cross-sectional biological age differences are not equivalent to longitudinal aging rates, but they may still encode structured information about the direction of methylome change along an aging-related axis.

The use of biological age as pseudo-time is conceptually related to pseudotime approaches in single-cell transcriptomics, where cross-sectional molecular profiles are ordered to infer dynamic processes [8]. However, there are important differences. In single-cell studies, cells may represent transitional states within a developmental or differentiation process, whereas individuals in an aging cohort differ in genetics, environment, lifestyle, sex, cellular composition, disease status, and technical factors. Therefore, the inferred derivative should be interpreted as a population-level methylome tendency along a biological-age axis, not as a deterministic prediction of each individual’s future methylome.

The strongest result of this study is internal reproducibility: the derived PACE-like score was recoverable from held-out buccal methylomes. This suggests that the proposed score captures a coherent methylation pattern rather than random numerical variation. Sensitivity analyses further indicated that performance was reasonably stable across model configurations, particularly for Ridge regression with intermediate dynamic CpG counts. This supports the idea that the score reflects broad methylome structure rather than dependence on a single CpG subset.

A second important finding is that the buccal PACE-like score showed only modest association with measured DunedinPACE. This was expected to some degree. DunedinPACE is a blood-derived biomarker trained to approximate longitudinal physiological decline in the Dunedin Study [7], whereas our score was derived from buccal mucosa methylation and from a biological-age pseudo-time axis. The two scores therefore differ in tissue, training target, mathematical construction, and biological interpretation. Our results suggest that the proposed buccal PACE-like score should not be presented as a buccal implementation of DunedinPACE. Rather, it should be interpreted as a tissue-specific buccal methylome dynamics index.

The alternative DunedinPACE reconstruction analyses reinforce this conclusion. Direct supervised prediction of DunedinPACE from buccal methylomes showed weak performance, and imputing blood methylation values at DunedinPACE-related CpGs from buccal methylomes did not substantially improve sample-level DunedinPACE prediction. Interestingly, pooled CpG-level imputation showed very high correlation between observed and predicted blood beta-values, but this did not translate into strong DunedinPACE reconstruction. This illustrates an important point: recovering average CpG methylation patterns across CpGs is not equivalent to recovering a weighted composite aging biomarker within individuals.

From a biological perspective, buccal mucosa and blood differ substantially in cellular composition and environmental exposure. Blood methylation biomarkers may capture immune-cell composition, systemic inflammation, smoking exposure, and hematopoietic aging, whereas buccal mucosa may capture epithelial aging, oral environmental exposure, local inflammation, and oral epithelial turnover. These tissue differences may explain why a buccal PACE-like score is only modestly correlated with blood DunedinPACE. Rather than being a weakness, this may indicate that buccal methylome dynamics represent a complementary tissue-specific dimension of aging.

The proposed framework has several limitations. First, the data are cross-sectional, and the inferred velocity is not directly observed longitudinal change. Second, the method is dependent on the biological age estimator used as pseudo-time. A different biological age model may produce a different derivative field and therefore a different PACE-like score. Third, the same methylome data contribute both to pseudo-time ordering and to future-state projection, introducing potential circularity. Fourth, the sample size is modest relative to the dimensionality of the methylome, requiring strong regularization and cross-fitting. Fifth, cell composition, smoking, oral inflammation, and batch effects may influence buccal methylation and should be addressed more comprehensively in future studies. Finally, external validation and repeated sampling are needed before the score can be interpreted as a stable individual-level biomarker.

Future work should prioritize longitudinal buccal methylome datasets, short-term technical and biological repeatability studies, and associations with clinical phenotypes. Repeated buccal sampling would allow estimation of test-retest reliability, intra-individual variability, and minimal detectable change. Longitudinal follow-up would allow testing whether baseline buccal PACE-like scores predict actual future methylation change, clinical aging outcomes, or response to lifestyle interventions. In addition, integrating cell composition estimates, smoking methylation scores, and oral health markers may improve biological interpretability.

In summary, we present a pseudo-longitudinal derivative framework for estimating a buccal mucosa-derived PACE-like aging-rate score from cross-sectional methylome data. The score was reproducibly predicted from buccal methylomes but was not equivalent to blood-derived DunedinPACE. These findings support the feasibility of tissue-specific methylome dynamics modeling and provide a methodological basis for future longitudinal validation of buccal aging-rate biomarkers.

## Author Contributions

Tatsuma Shoji (TS) performed the data analysis, generated all figures and tables, and drafted the manuscript. Ryo Nakaki (RN) led the project’s overall direction and management.

## Data Availability Statement

The data presented in this study are available from the corresponding author upon reasonable request, subject to institutional and ethical restrictions. DNA methylation data may contain potentially identifiable genomic information and therefore are not publicly released without appropriate data access approval.

## Acknowledgments

We thank Editage (www.editage.com) for English language editing.

## References

1. Horvath S. DNA methylation age of human tissues and cell types. Genome Biol. 2013;14(10):R115. doi:10.1186/gb-2013-14-10-r115.

2. Hannum G, Guinney J, Zhao L, Zhang L, Hughes G, Sadda S, Klotzle B, Bibikova M, Fan JB, Gao Y, Deconde R, Chen M, Rajapakse I, Friend S, Ideker T, Zhang K. Genome-wide methylation profiles reveal quantitative views of human aging rates. Mol Cell. 2013;49(2):359–367. doi:10.1016/j.molcel.2012.10.016.

3. Levine ME, Lu AT, Quach A, Chen BH, Assimes TL, Bandinelli S, Hou L, Baccarelli AA, Stewart JD, Li Y, Whitsel EA, Wilson JG, Reiner AP, Aviv A, Lohman K, Liu Y, Ferrucci L, Horvath S. An epigenetic biomarker of aging for lifespan and healthspan. Aging (Albany NY). 2018;10(4):573–591. doi:10.18632/aging.101414.

4. Lu AT, Quach A, Wilson JG, Reiner AP, Aviv A, Raj K, Hou L, Baccarelli AA, Li Y, Stewart JD, Whitsel EA, Assimes TL, Ferrucci L, Horvath S. DNA methylation GrimAge strongly predicts lifespan and healthspan. Aging (Albany NY). 2019;11(2):303–327. doi:10.18632/aging.101684.

5. Horvath S, Raj K. DNA methylation-based biomarkers and the epigenetic clock theory of ageing. Nat Rev Genet. 2018;19(6):371–384. doi:10.1038/s41576-018-0004-3.

6. Bell CG, Lowe R, Adams PD, Baccarelli AA, Beck S, Bell JT, Christensen BC, Gladyshev VN, Heijmans BT, Horvath S, Ideker T, Issa JPJ, Kelsey KT, Marioni RE, Reik W, Relton CL, Schalkwyk LC, Teschendorff AE, Wagner W, Zhang K, Rakyan VK. DNA methylation aging clocks: challenges and recommendations. Genome Biol. 2019;20(1):249. doi:10.1186/s13059-019-1824-y.

7. Belsky DW, Caspi A, Corcoran DL, Sugden K, Poulton R, Arseneault L, Baccarelli A, Chamarti K, Gao X, Hannon E, Harrington HL, Houts R, Kothari M, Kwon D, Mill J, Schwartz J, Vokonas P, Wang C, Williams BS, Moffitt TE. DunedinPACE, a DNA methylation biomarker of the pace of aging. eLife. 2022;11:e73420. doi:10.7554/eLife.73420.

8. Trapnell C, Cacchiarelli D, Grimsby J, Pokharel P, Li S, Morse M, Lennon NJ, Livak KJ, Mikkelsen TS, Rinn JL. The dynamics and regulators of cell fate decisions are revealed by pseudotemporal ordering of single cells. Nat Biotechnol. 2014;32(4):381–386. doi:10.1038/nbt.2859.

9. Du P, Zhang X, Huang CC, Jafari N, Kibbe WA, Hou L, Lin SM. Comparison of beta-value and M-value methods for quantifying methylation levels by microarray analysis. BMC Bioinformatics. 2010;11:587. doi:10.1186/1471-2105-11-587.

10. Shoji T, Tomo Y, Nakaki R. Prediction of biological age and blood biomarkers from DNA methylation profiles measured by the Methylation Screening Array: development and validation of models on Japanese data [Preprint]. bioRxiv. 2026. 10.64898/2026.02.06.703638.

11. Triche TJ Jr, Weisenberger DJ, Van Den Berg D, Laird PW, Siegmund KD. Low-level processing of Illumina Infinium DNA Methylation BeadArrays. Nucleic Acids Res. 2013;41:e90. doi:10.1093/nar/gkt090.

12. Zhou W, Triche TJ Jr, Laird PW, Shen H. SeSAMe: reducing artifactual detection of DNA methylation by Infinium BeadChips in genomic deletions. Nucleic Acids Res. 2018;46:e123. doi:10.1093/nar/gky691.

13. Aryee MJ, Jaffe AE, Corrada-Bravo H, Ladd-Acosta C, Feinberg AP, Hansen KD, et al. Minfi: a flexible and comprehensive Bioconductor package for the analysis of Infinium DNA methylation microarrays. Bioinformatics. 2014;30:1363–9. doi:10.1093/bioinformatics/btu049.

14. Shoji T, Tomo Y, Nakaki R. Predicting Biological Age and Clinical Biomarkers from DNA Methylation Profiles of Cheek Mucosa [Preprint]. bioRxiv. 2026 May 14. doi:10.64898/2026.05.12.724485.

